# Effects of targeted assistance and perturbations on the relationship between pelvis motion and step width in people with chronic stroke

**DOI:** 10.1101/2020.05.06.080705

**Authors:** Nicholas K. Reimold, Holly A. Knapp, Alyssa N. Chesnutt, Alexa Agne, Jesse C. Dean

## Abstract

**Background:** People with chronic stroke (PwCS) often exhibit a weakened relationship between pelvis motion and paretic step width, a behavior important for gait stabilization. We have developed a force-field able to manipulate this relationship on a step-by-step basis.

**Objective:** The objective of this study was to investigate the effects of a single exposure to our novel force-field on the step-by-step modulation of paretic step width among PwCS, quantified by the partial correlation between mediolateral pelvis displacement at the start of a step and paretic step width (step start paretic ρ_disp_).

**Methods:** Following a 3-minute period of normal walking, participants were exposed to 5-minutes of either force-field assistance (n=10; pushing the swing leg toward a mechanically-appropriate step width) or perturbations (n=10; pushing the swing leg away from a mechanically-appropriate step width). This period of assistance or perturbations was followed by a 1-minute catch period to identify any after-effects, a sign of sensorimotor adaptation.

**Results:** We found that assistance did not have a significant direct effect or after-effect on step start paretic ρ_disp_. In contrast, perturbations directly reduced step start paretic ρ_disp_ (p=0.004), but were followed by an after-effect in which this metric was increased above the baseline level (p=0.02).

**Conclusions:** These initial results suggest that PwCS have the ability to strengthen the link between pelvis motion and paretic foot placement if exposed to a novel mechanical environment, which may benefit gait stability. Future work is needed to determine whether this effect can be extended with repeated exposure to force-field perturbations.

## Introduction

Community-dwelling people with chronic stroke (PwCS) have an elevated fall-risk, with falls most commonly occurring during walking.^1^ Many falls are not due to external perturbations (e.g. slips or trips), but are instead attributed to “intrinsic factors” such as impaired balance.^2^ Balance deficits have been assessed using metrics of varying complexity, generally finding that relative to age-matched controls, PwCS tend to walk with: shorter and wider steps; more variable step lengths, times, and widths; more variable mediolateral and anteroposterior margins of stability; and larger mediolateral and vertical local divergence exponents (measuring the response to small perturbations).^3-5^ Several of these metrics can prospectively predict falls among PwCS (i.e. decreased step length, increased step time and length variability, increased local divergence exponent for mediolateral sacrum motion).^6^ Mediolateral balance deficits are a particular focus of attention, as most falls among PwCS occur sideways.^7^

Biomechanically, appropriate mediolateral foot placement is an important mechanism for ensuring walking balance.^8^ Neurologically-intact controls tend to place their swing foot relatively lateral for steps in which the pelvis has a large mediolateral displacement or velocity away from the stance foot, and relatively medial when the pelvis remains close to the stance foot. This behavior reduces the risk of the center of mass (CoM) moving lateral to the base of support, and can be quantified by relating fluctuations in pelvis motion to step-by-step adjustments in step width.^9^ Importantly, pelvis motion at the start of a step predicts the upcoming step width, likely due to a combination of passive body mechanics and active control evidenced by modulation of swing phase gluteus medius activity.^10^ In PwCS, the relationship between pelvis motion and step width is weaker for paretic steps, quantified using the partial correlation between mediolateral pelvis displacement and step width (ρ_disp_).^11^ This paretic side deficit is likely due in part to altered active control, as the normal link between pelvis motion and paretic swing phase gluteus medius activity is disrupted in PwCS with poor balance.^12^

Given the post-stroke deficits in the foot placement gait stabilization strategy, we have developed a novel force-field to manipulate the relationship between pelvis motion and step width.^13-14^ In neurologically-intact controls, we have used this force-field to *assist* appropriate mediolateral foot placement, encouraging wide steps when the pelvis is displaced far from the stance foot and narrow steps when the pelvis is close to the stance foot. This approach is based on the proposal that assisting a movement pattern can strengthen an individual’s ability to perform this movement independently, due in part to enhanced sensory feedback.^15-16^ Experimentally, we found that assistance had the intended direct effect of increasing ρ_disp_, indicating a stronger correlational link between pelvis displacement and step width.^14^ Conversely, we also used the force-field to *perturb* appropriate foot placement (e.g. encouraging narrow steps when a wide step is warranted), based on the principle that increasing movement errors can promote motor adaptation and learning.^17^ Here, we found that force-field perturbations directly decreased ρ_disp_, weakening the link between pelvis displacement and step width.^14^ As our ultimate goal is to alter gait behavior outside of the force-field, we quantified whether exposure to force-field assistance or perturbations also produced after-effects in ρ_disp_ – an indicator of altered sensorimotor control.^17^ Such after-effects were observed, as assistance was followed by short-lived decreases in ρ_disp_, while perturbations were followed by short-lived increases in ρ_disp_.^14^

While our initial results in neurologically-intact controls are promising, it is unclear whether the relationship between pelvis motion and step width can be similarly altered in PwCS. Encouragingly, prior work has found that novel mechanical environments can indeed evoke changes in gait behavior among this population. Most notably, using a split-belt treadmill to amplify step length asymmetries can cause after-effects in which these asymmetries are temporarily reduced.^18^ Similarly, unilaterally resisting forward leg swing can produce after-effects with reduced step length asymmetry.^19-20^ In the frontal plane, forces pushing the pelvis toward the paretic side can cause after-effects in which the pelvis stays displaced farther away from the paretic leg^21^ – an effect that may not be deemed beneficial. In each case, a novel mechanical environment produced changes in a targeted biomechanical behavior.

The purpose of this study was to investigate the effects of a single force-field exposure on the step-by-step modulation of paretic step width among PwCS. Specifically, we tested whether applying force-field assistance or perturbations had direct effects on the relationship between paretic step width and mediolateral pelvis displacement at the start of the step, as well as whether after-effects were observed once these forces ceased. Based on our prior results, we hypothesized that assistance would directly strengthen the relationship between pelvis displacement and step width (increase ρ_disp_), but would be followed by negative after-effects as ρ_disp_ would decrease relative to baseline once the assistance ceased. Conversely, we hypothesized that perturbations would have the direct effect of decreasing ρ_disp_, but would be followed by positive after-effects in which ρ_disp_ increased. Such results would demonstrate that PwCS have the ability to modulate the sensorimotor control underlying the foot placement stabilization strategy.

## Methods

### Participants

Twenty PwCS participated in this study, a sample size based on preliminary results presented in the Appendix. Basic information regarding participant demographics and function is presented in Table 1. Participants were recruited from a previous screening session used to determine whether they met the study’s inclusion and exclusion criteria. Inclusion criteria were: age≥21 years; experience of a stroke 6 months prior; residual paresis in a lower extremity; preferred overground gait speed of at least 0.2 m/s; ability to walk for 3-minutes without a cane or walker; a step start paretic ρ_disp_ value (described in the Introduction and detailed below) of below 0.56, the lower bound of the 95% confidence interval for this metric among neurologically-intact participants walking at a typical preferred speed.^22^ Exclusion criteria were: a resting heart rate above 110 beats/min or blood pressure above 200/110 mm Hg; history of cardiac dysfunction; pre-existing neurological disorders; severe visual impairment; history of pulmonary embolism; uncontrolled diabetes; orthopedic injuries or conditions with the potential to alter foot placement. All participants provided written informed consent using a form approved by the Medical University of South Carolina Institutional Review Board and consistent with the Declaration of Helsinki.

**Table 1.**
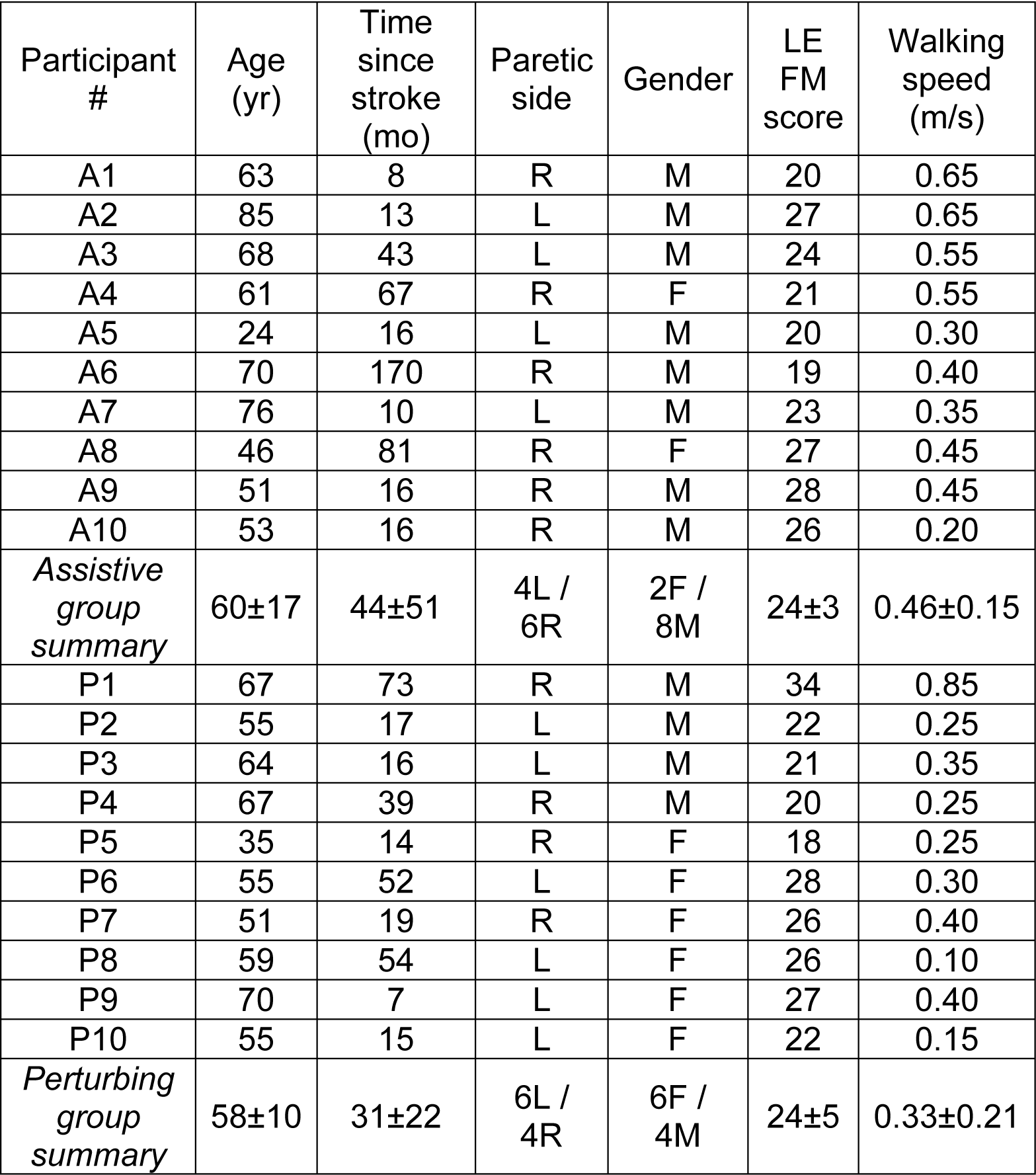
Participant demographic characteristics, lower extremity Fugl-Meyer (LE FM) motor scores, and treadmill walking speeds. Numerical summary scores are presented as means ± standard deviations.

### Experimental Protocol

Participants were randomly assigned to the Assistive (n=10) or Perturbing (n=10) group, and blinded to their group assignment. Participants in both groups performed a series of 3-minute walking trials at their self-selected speed, defined as the speed used “to walk around your house or the store” and identified by iteratively increasing the treadmill speed. All participants wore a harness attached to an overhead rail that did not support body weight, but would have prevented a fall in case of a loss of balance. Participants were not permitted to hold onto a handrail, which can alter the control of walking balance.^23^

For the first treadmill trial (Normal), participants were not interfaced with the force-field. For the second trial, the force-field remained in either Assistive or Perturbing mode (described below) throughout the trial, corresponding to the participant’s group assignment. For the third trial, the force-field was in Assistive/Perturbing mode for the first 2-minutes, before switching to Transparent mode (described below) for the final minute. Our prior work in neurologically-intact participants found that 5-minutes of force-field exposure was sufficient to produce after-effects during the subsequent minute of walking [Heitkamp et al. 2019; Reimold et al. 2019]. Here, trial duration was 3-minutes to reduce the risk of fatigue.

### Force-field Design and Control

The force-field used to exert mediolateral forces on participants’ legs has been described in detail previously.^14,24^ Briefly, forces are applied using steel wires running parallel with the treadmill belts and in series with extension springs. The wires pass through cuffs worn on the lateral side of the shanks, allowing free anteroposterior and vertical leg motion. Linear actuators (UltraMotion; Cutchoge, NY, USA) are used to vary the mediolateral position of the wires interfacing with the swing leg, encouraging participants to step to a targeted step width. The mediolateral leg forces are proportional to the leg’s deviation from the targeted mediolateral location, with an effective mediolateral stiffness of 180 N/m. The force-field was controlled using feedback from active LED markers (PhaseSpace; San Leandro, CA, USA) placed on the participants’ sacrum, heels, and leg cuffs.

The Transparent mode was designed to minimize the mediolateral leg forces experienced by users, as the force-field wires were continuously repositioned to stay aligned with the corresponding leg cuff.

The Assistive mode was designed to assist participants with placing their swing foot in a mechanically-appropriate mediolateral location. For example, if the pelvis was far to the right of the stance foot at the start of a right step, the force-field would push the swing leg laterally to encourage a wide step. The targeted location was determined for each step based on the following equation, derived empirically from neurologically-intact participants:^22^

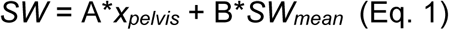

Here, x_pelvis_ is the mediolateral location of the sacrum relative to the stance heel at the start of the step. SW_mean_ is the participant’s mean step width, as calculated from the last 50-steps of the initial Normal walking trial. A and B are coefficients that vary with participant walking speed, as illustrated in Figure 1A. We chose this control equation as pilot experiments (Appendix) found it to be sufficient to produce measurable changes in the relationship between pelvis displacement and step width.

**Figure 1.**
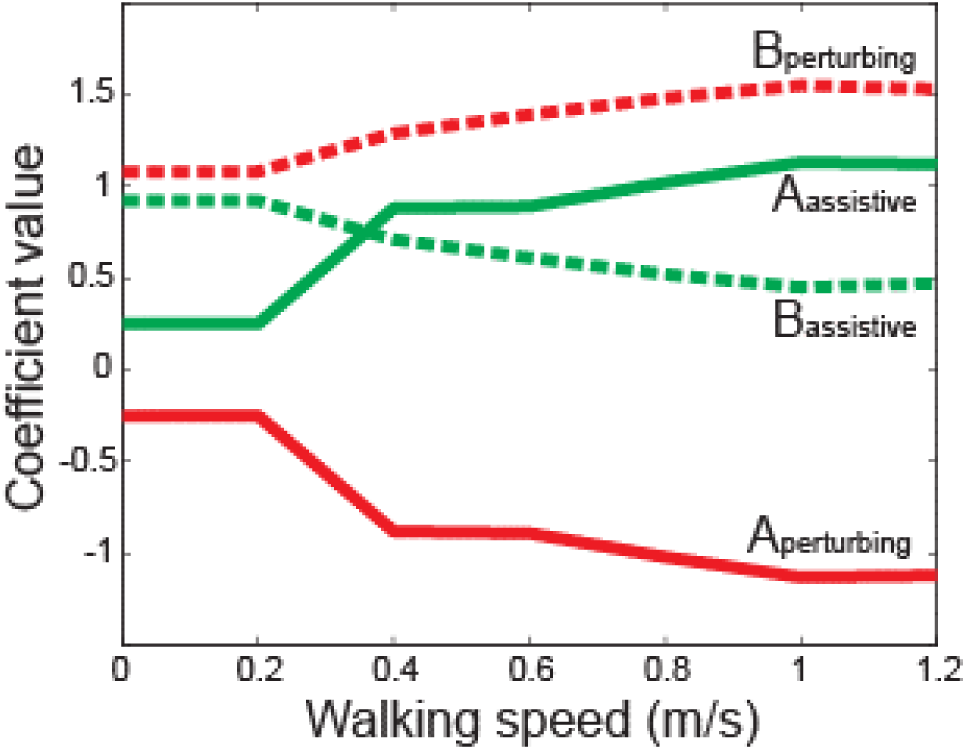
Force-field equation coefficient values (A & B) for the Assistive and Perturbing conditions, as labeled. These values vary with walking speed, based on empirically-derived best-fit coefficients from neurologically-intact control participants walking at a range of speeds (0.2, 0.4, 0.6, 0.8, 1.0, 1.2 m/s), and linearly interpolated between these values. For any speeds below 0.2 m/s, we used the 0.2 m/s coefficients.

The Perturbing mode was designed to push participants’ steps *away from* a mechanically-appropriate location, potentially requiring an active response to prevent a loss of balance. Here, if the pelvis was far to the right of a stance foot at the start of a right step, the force-field would push the swing leg medially to encourage a narrow step, increasing the risk of the center of mass moving lateral to the base of support during the subsequent stance phase. The force-field control equation took the same form as Equation 1 above, with coefficients illustrated in Figure 1B.

### Data Collection and Processing

The locations of LED markers placed on the sacrum, bilateral heels, and bilateral leg cuffs were sampled at 120 Hz, and low-pass filtered at 10 Hz. Each step start was defined to occur when the ipsilateral heel velocity changed from posterior to anterior.^25^ Each step end was defined to occur when the contralateral heel velocity changed from posterior to anterior. For each minute of walking we calculated several gait metrics focused on the step-by-step relationship between pelvis motion and step width, as well as several traditional gait metrics.

Our primary outcome measure was the partial correlation between the step width and the mediolateral displacement of the pelvis from the stance foot at the start of the paretic step, accounting for mediolateral velocity of the pelvis (hereby termed step start paretic ρ_disp_). This metric quantifies a relationship that is e important for mediolateral balance during walking,^8^ with a focus on potential adjustments that can occur during a step. We have previously shown that this metric is decreased for paretic steps among PwCS.^11^ While mediolateral pelvis velocity plays a secondary role in influencing step width,^22^ this relationship is not altered for paretic steps.^11^

Although our primary focus is on the link between paretic step width and pelvis displacement at the start of a step, for illustrative purposes we quantified this relationship for pelvis displacement throughout the course of both paretic and non-paretic steps. Specifically, at each normalized time point in a step (from 0-100), we calculated ρ_disp_ based on the pelvis displacement and velocity values at this time point. The value of this metric at the end of a step (step end ρ_disp_) is of secondary interest, as it is conceptually similar to step-by-step variability in mediolateral margin of stability values calculated from extrapolated center of mass^9^ – a more common metric of walking balance.

We defined step width as the mediolateral displacement between the ipsilateral heel marker at the step end and the contralateral heel marker at the step start. We defined mediolateral foot placement as the mediolateral displacement between the sacrum and the ipsilateral heel at the step end. Finally, we defined step length as the difference between the anterior position of the ipsilateral heel at the step end and the anterior position of the contralateral heel at the previous step end, accounting for treadmill speed. We calculated the mean of these metrics for both paretic and non-paretic steps.

### Statistics

Due to the relatively small sample size, we used non-parametric statistics to investigate potential direct effects and after-effects of force-field exposure. For our primary analysis in the Assistive group, we applied Wilcoxon signed rank tests to compare the average value of step start paretic ρ_disp_ between the Normal and Assistive periods (direct effects), and to compare this metric between the Normal and Transparent periods (after-effects). To account for performing two comparisons, the alpha value indicating significance was set to 0.025. This primary analysis was repeated for the Perturbing group to detect direct effects and after-effects on step start paretic ρ_disp_. In secondary analyses, an identical statistical approach was applied for both the Assistive and Perturbing groups to test for direct effects and after-effects on: step end paretic ρ_disp_; step start non-paretic ρ_disp_; step end non-paretic ρ_disp_; mean step width; mean step length; mean mediolateral foot placement. Each of the traditional gait metrics was compared for both paretic and non-paretic steps.

## Results

The partial correlation between pelvis displacement and step width (ρ_disp_) increased over the course of a step, as illustrated for participants in both the Assistive (Fig. 2A-B) and Perturbing (Fig. 2C-D) groups. The magnitude of ρ_disp_ during a step was generally smaller for paretic steps than non-paretic steps, and the force-field effects differed between the Assistive and Perturbing modes.

**Figure 2.**
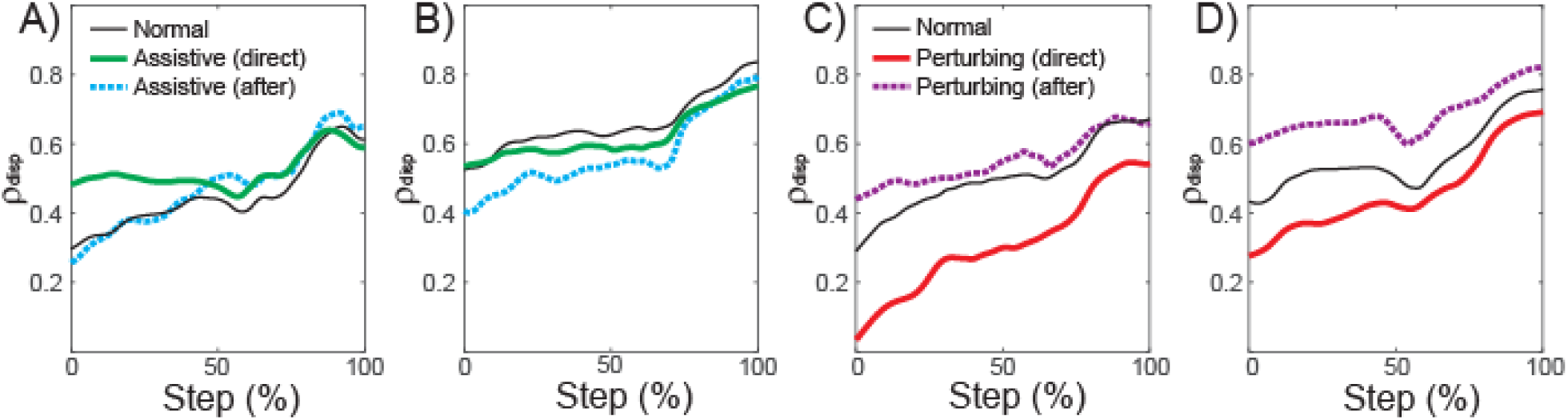
ρ_disp_ values calculated throughout the step, from step start (0%) to step end (100%). The direct effects and after-effects of assistance are illustrated for steps taken with the paretic leg (A) and the non-paretic leg (B). Similarly, the direct effects and after-effects of perturbations are illustrated for paretic steps (C) and non-paretic steps (D). Plots illustrate the mean value of these metrics across participants. Intersubject variability is not depicted due to excessive overlap, but is apparent from Figures 3-4 illustrating individual values.

Changes in paretic ρ_disp_ were more apparent with perturbations than assistance. Assistance did not have a significant direct effect (p=0.037; Fig. 3A) on our primary metric of step start paretic ρ_disp_. Nor were any significant after-effects (p=0.77; Fig. 3B) observed once the assistance ceased. In contrast, perturbations directly caused a significant decrease (p=0.004; Fig. 3C) in step start paretic ρ_disp_, followed by a significant increase (p=0.02; Fig. 3D) after the perturbations ceased. For our secondary measure of step end paretic ρ_disp_, assistance had no significant direct (p=0.56; Fig. 3E) or after effects (p=0.38; Fig. 3F). Perturbations significantly decreased step end paretic ρ_disp_ while applied (p=0.014; Fig. 3G), but had no significant after-effects (p=0.85; Fig. 3H).

**Figure 3.**
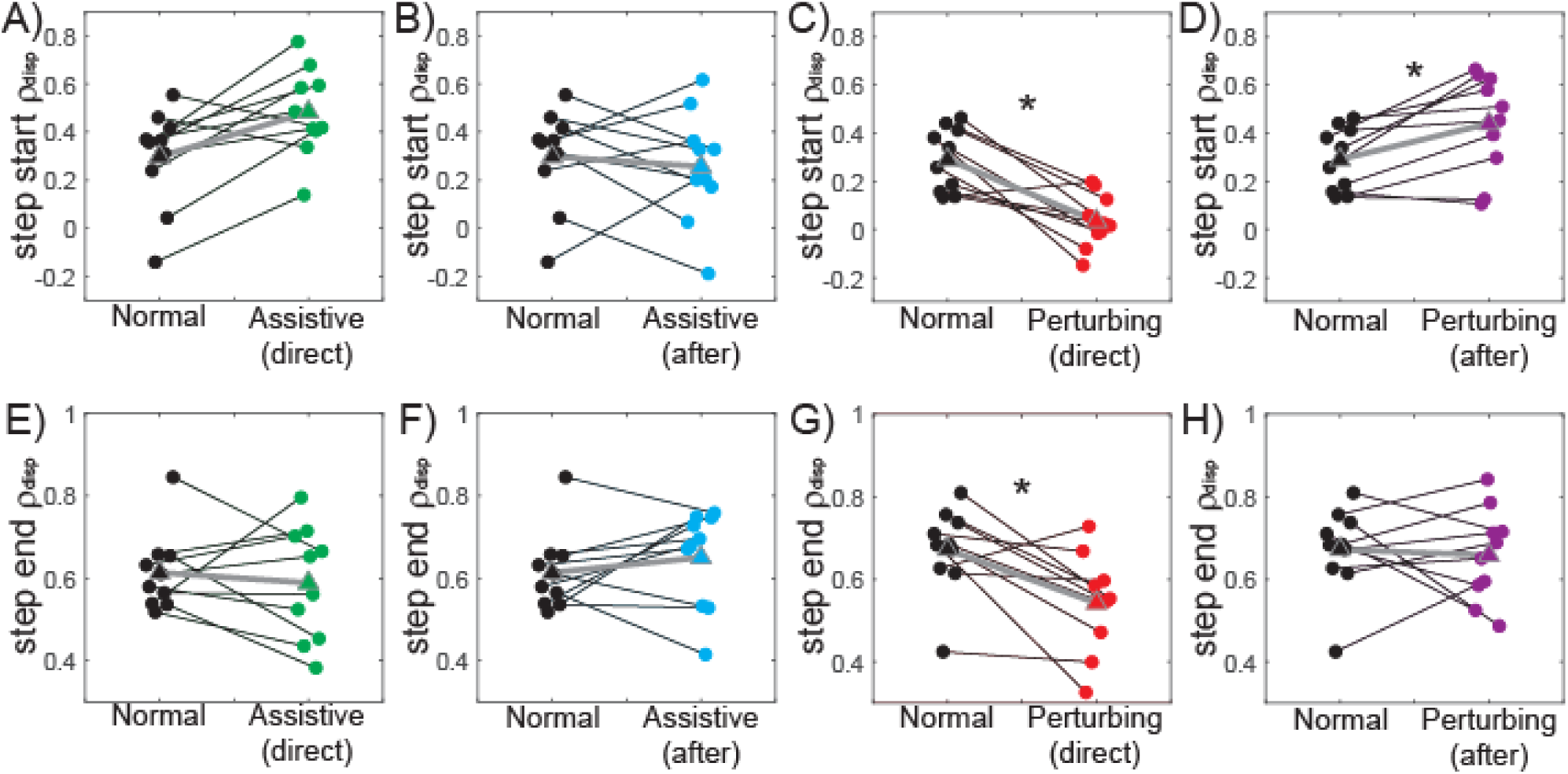
Force-field effects on paretic ρ_disp_. All panels illustrate individual participants’ values for these metrics, and the change relative to Normal walking. The top row (panels A-D) illustrates step start paretic ρ_disp_, while the bottom row (panels E-H) illustrates step end paretic ρ_disp_. From left to right, the panels in each row illustrate the direct effects of assistance, the after-effects of assistance, the direct effects of perturbations, and the after-effects of perturbations. Small circles and thin black lines indicate individual values, outlined triangles and thick gray lines indicate group means, and statistical significance is illustrated with asterisks (*). Individual data points are slightly offset on the x-axis to reduce overlap.

The force-field’s effects on non-paretic steps differed somewhat from paretic steps. Assistance again had no significant direct effects (p=0.92; Fig. 4A) or after-effects (p=0.11; Fig. 4B) on step start non-paretic ρ_disp_. While perturbations directly caused a significant decrease in step start non-paretic ρ_disp_ (p=0.01; Fig. 4C), no significant after-effects (p=0.11; Fig. 4D) were observed. Both assistance and perturbations appeared to directly reduce step end non-paretic ρ_disp_ (Fig. 4E, G), although neither of these reductions reached significance (Assistive p=0.037; Perturbing p=0.038). Neither assistance (p=0.084; Fig. 4F) nor perturbations (p=0.13; Fig. 4H) caused significant after-effects on step end non-paretic ρ_disp_.

**Figure 4.**
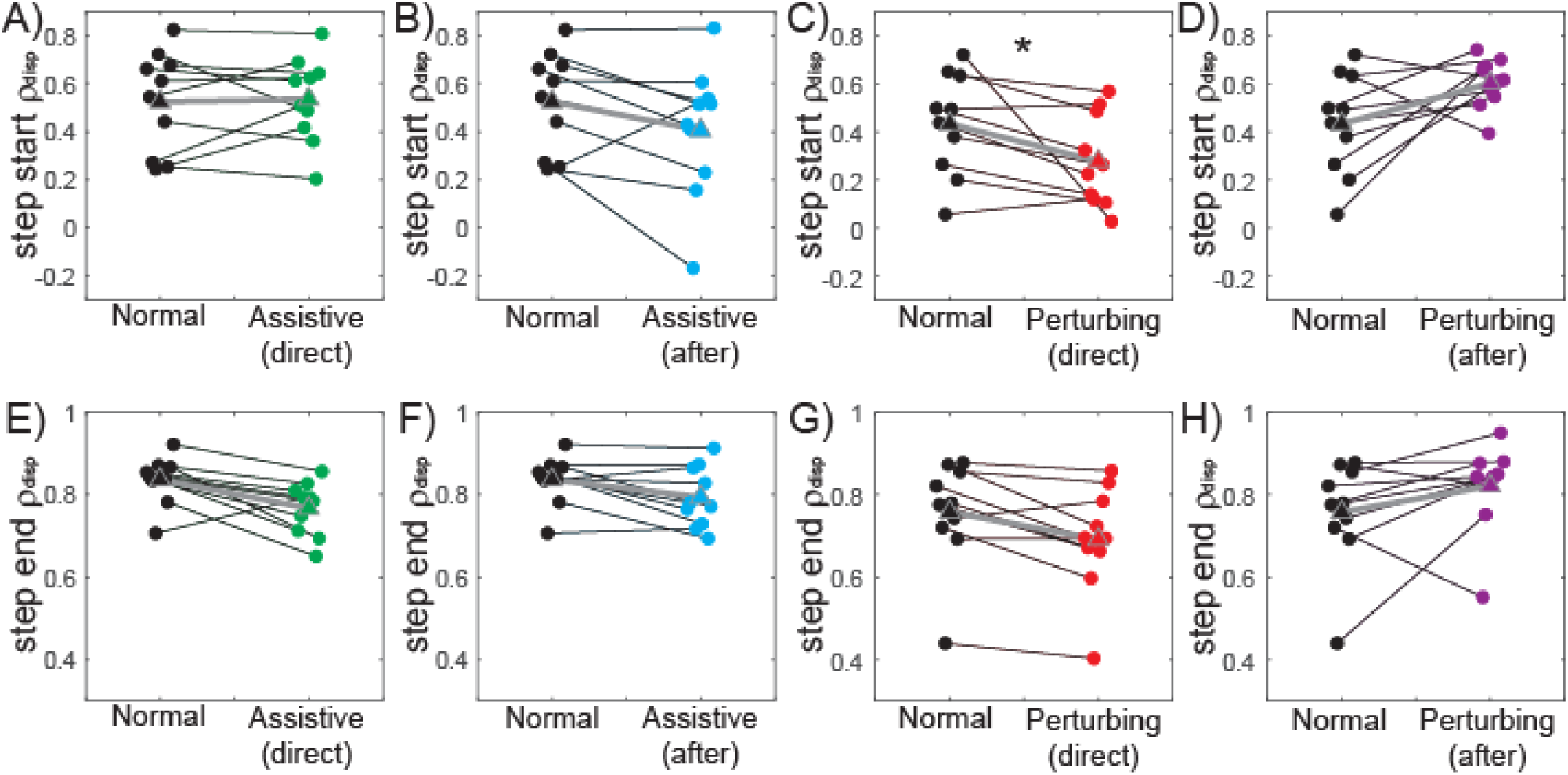
Force-field effects on non-paretic ρ_disp_. The figure structure matches that described for Figure 3.

The effects of assistance and perturbations on traditional gait metrics are presented in Table 2. Assistance had no significant effect on step width, step length, or mediolateral foot placement. In contrast, perturbations had the direct effect of increasing step width (due to more lateral paretic foot placement), but did not influence step length. No after-effects in these metrics were present once the perturbations ceased.

**Table 2.** Effects of assistance and perturbations on gait spatial measures, all in units of mm. Values are presented as means ± standard deviations, followed by the corresponding p-value for comparisons with the baseline Normal condition. Statistically significant comparisons (p<0.025) are indicated using bolded italics.

|  | Assistive group |  |  | Perturbing group |  |  |
| --- | --- | --- | --- | --- | --- | --- |
|  | Normal | Assistive direct effects | Assistive after-effects | Normal | Perturbing direct effects | Perturbing after-effects |
| Paretic SW | 205±40 | 206±46<br>(p=0.92) | 199±34<br>(p=0.084) | 162±64 | **176±66**<br>**(p=0.014)** | 169±62<br>(p=0.49) |
| Paretic SL | 368±134 | 366±121<br>(p=0.92) | 369±123<br>(p=1) | 245±107 | 239±119<br>(p=0.23) | 241±125<br>(p=0.56) |
| Paretic FP | 86±19 | 91±20<br>(p=0.32) | 88±22<br>(p=0.49) | 81±44 | **92±51**<br>**(p=0.002)** | 86±45<br>(p=0.23) |
| Non-paretic SW | 204±40 | 206±46<br>(p=1) | 199±34<br>(p=0.084) | 162±64 | **176±66**<br>**(p=0.014)** | 169±62<br>(p=0.49) |
| Non-Paretic SL | 361±102 | 355±102<br>(p=0.28) | 369±109<br>(p=0.63) | 225±133 | 223±128<br>(p=0.77) | 229±131<br>(p=0.28) |
| Non-Paretic FP | 55±33 | 50±32<br>(p=1) | 49±30<br>(p=0.56) | 29±41 | 33±44<br>(p=0.13) | 34±47<br>(p=0.19) |

## Discussion

We investigated whether a single exposure to assistive or perturbing forces influenced the relationship between pelvis motion and step width in PwCS, with results partially supporting our hypotheses. Force-field assistance did not consistently strengthen the relationship between pelvis displacement at the start of a step and paretic step width, and had no notable after-effects. Conversely, force-field perturbations directly weakened this relationship, as the stepping leg was pushed away from a mechanically-appropriate location. Perturbation cessation was followed by a period of after-effects in which the relationship between pelvis displacement at the start of a step and step width was strengthened relative to baseline, indicative of altered sensorimotor control.

Force-field assistance did not have a consistent effect on the relationship between pelvis displacement and paretic step width. Unlike in neurologically-intact participants,^14^ the direct effects of assistance on step start paretic ρ_disp_ did not reach our set level of significance (p=0.038). However, we note that only the two participants with the highest baseline values of step start paretic ρ_disp_ did not exhibit a clear increase with assistance (see Fig 3A). Additionally, assistance did significantly increase step start paretic ρ_disp_ for the separate population of PwCS in the pilot experiment described in the Appendix. Perhaps for the participants with near-normal paretic ρ_disp_ values, assistive forces interfered with the baseline gait pattern perceived as stable, and thus were resisted. Similar counter-intuitive effects of assistance have been previously observed in PwCS, with artificial plantarflexor power reducing anterior propulsive forces,^26^ and knee flexion assistance for swing phase ground clearance increasing leg circumduction.^27^ Beneficial effects may require adjustment of the force-field control paradigm, such that some participants do not perceive the assistance as perturbations that should be resisted. Beyond the equivocal direct effects, we observed no consistent after-effects on step start paretic ρ_disp_ following force-field assistance. These initial results thus provide no evidence that a brief period of assistance allows PwCS to discover and maintain the targeted gait pattern, although this general approach is common in clinical practice.^28^

In contrast to force-field assistance, perturbations had significant direct effects and after-effects on the relationship between pelvis displacement at the start of a step and paretic step width. These effects were similar to those in neurologically-intact controls;^14^ perturbations directly caused a decrease in step start paretic ρ_disp_, but a subsequent positive after-effect once the perturbations ceased. This after-effect demonstrates that as a group, PwCS have the ability to walk with a stronger relationship between pelvis motion and paretic step width – a relationship often weakened in this population.^11^ As with assistance, visual inspection of our results (Fig. 3D) suggests a possible effect of baseline function. The only participants not to exhibit a positive after-effect were the two with the lowest baseline value of step start paretic ρ_disp_. Previous work in sensorimotor adaptation and learning has suggested just such an effect – individuals with a low skill level may not benefit from perturbations that augment movement errors.^16,29^ Instead, task difficulty should be based on an individual’s skill level, an idea formalized in the challenge point framework.^30^

As with our primary metric (paretic step start ρ_disp_), our secondary metrics were more clearly affected by perturbations than assistance. Specifically, assistance did not significantly strengthen the relationship between pelvis displacement and step width for non-paretic steps, or when calculated based on pelvis displacement at the end of the step. In fact, we observed a non-significant trend (p=0.038) for assistance to reduce non-paretic step end ρ_disp_, contrary to our expectations. This decrease may be due to the preference of many PwCS to keep their non-paretic leg close to their CoM;^11,12,31^ assistive forces that pushed the non-paretic leg laterally may have been perceived as de-stabilizing, and resisted. Assistance also did not produce consistent after-effects, and did not influence mean step width, step length, or foot placement. On the other hand, perturbations directly reduced ρ_disp_ magnitude for both legs and both tested time points (although this did not reach significance for step end non-paretic ρ_disp_; p=0.037), indicating effective perturbation of the targeted stabilization strategy. Following the perturbations, we did not observe significant positive after-effects for our secondary ρ_disp_ measures. Based on our recent work in neurologically-intact participants, the sensorimotor adjustments that underlie potential after-effects may require longer than 5-minutes of perturbation exposure.^24^ Alternatively, perhaps sensorimotor adjustments could be facilitated by switching between mechanical environments more often (e.g. more brief periods in Transparent mode), as can accelerate relearning during split-belt walking.^32^ In addition to altering the relationship between pelvis motion and step width, applying perturbations directly caused an increase in step width and more lateral paretic foot placement. Importantly, the equation used to control the perturbations accounted for each participant’s mean step width, and thus did not directly push the legs to take wider steps. Instead, the wider steps and more lateral paretic foot placement is likely due to perceived instability, particularly when transitioning into paretic stance.^10,33^

Our primary finding is that as a group, PwCS can adjust their gait pattern so pelvis displacement at the start of a step is more closely linked to paretic step width – thus strengthening a presumably useful gait stabilization strategy. Our presumption that this strategy is useful is based on the consistent theme from model simulations,^34^ bipedal robot design,^35^ and assistive exoskeletons^36-37^ that adjusting mediolateral foot placement is an efficient method of ensuring walking balance. However, the finding that many PwCS have the ability to strengthen this strategy after a short period of perturbations raises the question of why they do not simply do so in their baseline gait pattern. This use of a gait pattern perceived by observers as non-ideal is common post-stroke, perhaps best exemplified by step length asymmetry. Many PwCS walk with substantial step length asymmetries, but can adopt a more symmetric gait after a brief period of split-belt walking in which these asymmetries are amplified.^18^ One possible explanation for the post-stroke use of “non-ideal” movement patterns is that individuals develop and strengthen these habits in the acute or sub-acute phase, when brain plasticity is heightened.^38^ In the early stages after a stroke when control of the paretic leg can be quite impaired, a strategy of placing the paretic leg far laterally irrespective of pelvis motion could be beneficial by simplifying control, reducing the risk of a loss of balance toward the paretic side, and allowing gravity to redirect the CoM back toward the stronger leg.^39^ This habitual movement pattern may be retained even as paretic leg function improves. Alternatively, it is possible that walking with consistently lateral paretic foot placement (or notable step length asymmetries) has benefits in the chronic stage that are not yet known. The present study is not able to differentiate between these possibilities.

This initial investigation in PwCS focused on a single exposure to assistance or perturbations, and cannot provide insight into longer-term effects of walking in these novel mechanical environments. It is possible that participants gradually revert back to their baseline gait pattern once they return to the original mechanical environment (e.g. when perturbations cease).^17^ Indeed, while the present walking trials were brief to reduce the risk of fatigue, our prior work in neurologically-intact participants found that the positive after-effects following perturbations disappear in ∼1 minute of walking,^14^ but become longer lasting with repeated perturbation exposure.^24^ Other perturbation paradigms in patients with neurological injuries have similarly suggested the potential for sustained benefits with repeated exposure to a novel mechanical environment; step length asymmetry can be reduced with repeated periods of split-belt walking,^40^ step lengths can be increased with repeated swing leg resistance,^41^ and multiple bouts of pelvis perturbations can improve stabilizing responses.^42^ Speculatively, the evoked gait changes may only be sustained if they are perceived by the participant to be superior to their baseline movement pattern in some way.^17^ It is feasible that repeated experience with more mechanically-appropriate adjustments of step width will be perceived by PwCS as beneficial to their gait stability, allowing them to overcome habitual movement patterns. In an ongoing clinical trial, we are testing whether this occurs, as well as investigating links between changes in ρ_disp_ and other measures of negative balance outcomes (e.g. clinical tests of balance; fall incidence; fear of falling).

Our force-field is designed to perturb one aspect of gait – the relationship between pelvis motion and step width. While this behavior is believed to be important for gait stability,^8^ many other methods can be used perturb walking balance. Prior work in older adults has predominantly focused on translating a treadmill surface to elicit slip-like, trip-like, or mediolateral perturbations.^43-47^ More recently, similar approaches have been applied among PwCS,^48-51^ with results suggesting that this population can improve their balance responses. An alternative approach applies mediolateral forces to the trunk during walking. While these forces can assist symmetric weight shift by pushing the pelvis toward the paretic leg,^21,52-53^ other work has sought to either augment pelvis motion errors by pushing the pelvis farther toward the non-paretic leg,^52^ or apply unpredictable mediolateral forces.^42^ In the longer-term, it would be valuable to compare the effects of these various perturbation paradigms. Given the apparent specificity of perturbation responses,^44^ discrete perturbations requiring a reactive response (e.g. slip perturbations) may have a different effect on the sensorimotor control of balance than perturbations designed to encourage the restoration of a specific gait behavior (e.g. a stronger relationship between pelvis motion and step width).

A limitation of this study is the small sample size, which likely prevented us from detecting potential effects of force-field exposure with moderate effect sizes. However, the observed direct effects and after-effects of perturbations in this small population motivate more a detailed investigation. Additionally, the present work suggests that the baseline function of individual PwCS may influence the force-field’s effects (e.g. high functioning individuals did not exhibit a direct increase in step start paretic ρ_disp_ with assistance; low-functioning individuals did not exhibit a positive after-effect following perturbations). While similar effects of baseline function have previously been described,^16^ a substantially larger sample size would be required to investigate this topic rigorously.

## Conclusions

Targeted perturbations can cause people with chronic stroke to strengthen the relationship between pelvis motion and paretic step width, while assistance is generally less effective. Although this result suggests that many PwCS have the ability to successfully execute a gait stabilization strategy based on foot placement adjustments, future work is needed to achieve clinical relevance. Most notably, it is unclear whether the strengthened link between pelvis motion and paretic step width can be retained for longer periods of time (possibly with repeated exposure), whether changes in this gait behavior truly benefit walking balance in terms of clinical measures of function, and whether only a subset of participants will benefit from this approach.

## Acknowledgements

This work was supported by grants from the National Institutes of Health (1 R21 HD088869-01A1 and 1 P20 GM109040-01).

