## Appendix for "Effects of targeted assistance and perturbations on the relationship between pelvis motion and step width in people with chronic stroke"

### *Introduction*

People with chronic stroke (PwCS) often exhibit a weaker relationship between mediolateral pelvis displacement and step width for steps taken with their paretic leg [Stimpson et al. 2019]. As mechanics-dependent adjustments in step width are an important gait stabilization strategy [Bruijn and van Dieen 2018], this weaker relationship may contribute to deficits in post-stroke balance. To investigate this topic, developed a force-field able to encourage targeted step widths on a step-by-step basis [Nyberg et al. 2017].

In neurologically-intact controls, several biomechanical factors predict the step width for a given step, as influenced by both the body's passive dynamic properties and active control [Rankin et al. 2014]. Most prominently, people tend to place their swing leg more laterally to produce wider steps when the pelvis is displaced farther mediolaterally from the stance foot [Wang and Srinivasan 2014; Stimpson et al. 2018]. While not as strong a relationship, wider steps also tend to accompany larger mediolateral velocities away from the stance foot [Wang and Srinivasan 2014; Stimpson et al. 2018]. Additionally, these step-by-step fluctuations vary around a mean step width value that differs between individuals – likely due in part to differences in anthropometric factors such as pelvis width or leg circumference [Browning and Kram 2007], and possibly differences in an individuals' balance confidence [Hak et al. 2012].

Our prior work has investigated the effects of an assistive force-field that used various combinations of these three factors (pelvis displacement, pelvis velocity, mean step width) to predict mechanically-appropriate step widths in neurologically-intact participants [Heitkamp et al. 2019]. All force-field control equations included mediolateral pelvis displacement, and all equations significantly increased our primary outcome measure quantifying the relationship between pelvis displacement and step width – step start $\rho_{disp}$. These increases were significantly larger for equations that also included pelvis velocity, while the inclusion of mean step width appeared not to play an important role. However, it is unclear whether similar effects would be observed in PwCS, a population in whom step start $\rho_{disp}$ is likely reduced for steps taken with the paretic leg.

The purpose of this pilot experiment was to determine whether the relationship between pelvis displacement and paretic step width was differently affected by several candidate equations used to control force-field assistance. Specifically, we investigated the direct effects of equations that included various combinations of pelvis displacement, pelvis velocity, and the individual participant's mean step width. This experiment was not hypothesis-driven, but was used to identify the force-field equation to be used in our main experiment.

***Methods***

*Participants*

Twelve PwCS completed this study. Basic participant demographic information is provided in Table S1. The inclusion and exclusion criteria match those presented in the main text, with the exception that we did not have a maximum step start paretic $\rho_{disp}$ value for this preliminary experiment. All participants provided informed consent using a form approved by the Medical University of South Carolina Institutional Review Board and consistent with the Declaration of Helsinki.

*Experimental Protocol*

Participants performed a series of 2-minute walking trials at their normal walking speed, identified as described in the main text. For all trials, participants wore a harness to prevent falls and were not permitted to hold onto a handrail. For the first (Normal) treadmill trial, participants did not interact with the force field. Participants then performed four randomized-order walking trials in which assistance was controlled based on the four equations detailed below. The equations differed in terms of which combination of factors (pelvis displacement, pelvis velocity, and mean step width) were used to predict a mechanically-appropriate step width. These trials were separated by 2-minute wash-out periods in which participants walked with the force-field in Transparent mode. The present analyses focus only on the direct effects of the assistance, not the wash-out periods.

*Force-Field Control*

This preliminary experiment used the same force-field device described in the main text, and detailed in prior work [Nyberg et al. 2017; Heitkamp et al. 2019]. For each step, a mechanically-appropriate step width was calculated by one of four Assistive control equations taking the following form:

$$SW = \text{A}*x_{sacrum} + \text{B}*v_{sacrum} + \text{C}*SW_{mean} + \text{D} \qquad \text{(Eq. S1)}$$

Here, $x_{sacrum}$ is the mediolateral location of the sacrum relative to the stance heel at the start of the step. $v_{sacrum}$ is the mediolateral velocity of the sacrum at the start of the step. $SW_{mean}$ is the participant's mean step width, as calculated from the last 50-steps of the initial Normal walking trial. A, B, C, and D are coefficients that vary with participant walking speed, as illustrated in Figure S1. Note that for each equation, either the C or D coefficients are set to zero.

*Data Collection and Processing*

As in the main text, our primary outcome is step start paretic $\rho_{disp}$, as calculated from LED markers placed on the sacrum and bilateral heels, sampled at 120 Hz, and low-pass filtered at 10 Hz. Secondarily, we also report step start non-paretic $\rho_{disp}$, and step end $\rho_{disp}$ values for paretic and non-paretic steps.

*Statistics*

Due to the relatively small sample size in this experiment, we used non-parametric statistical comparisons. Specifically, we used Friedman's test ($\alpha$=0.05) to compare step start paretic $\rho_{disp}$ values between the baseline Normal walking condition and the four Assistive conditions (corresponding to four control equations). In the case of a significant effect, we used Tukey-Kramer post hoc tests to compare individual conditions. Secondarily, we repeated this analysis for our secondary measures of step start non-paretic $\rho_{disp}$, step end paretic $\rho_{disp}$, and step end non-paretic $\rho_{disp}$.

### *Results*

As observed in the main text, the partial correlation between pelvis displacement and step width ($\rho_{disp}$) increased over the course of a step, and was generally lower for paretic steps (Fig. S2A) than for non-paretic steps (Fig. S2B).

For paretic steps, step start $\rho_{disp}$ varied significantly (p=0.004; Fig. S3A) across the five conditions. Each of the assistance equations other than equation 1 caused a significant increase in step start paretic $\rho_{disp}$, and no significant differences were observed between equations. Step end paretic $\rho_{disp}$ did not differ significantly (p=0.082; Fig. S3B) across the five walking conditions. For non-paretic steps, no significant differences across the five walking conditions were observed for step start $\rho_{disp}$ (p=0.068; Fig. S3C) or step end $\rho_{disp}$ (p=0.87; Fig. S3D).

### *Discussion*

For our primary metric (step start paretic $\rho_{disp}$), we observed no significant differences across the four candidate equations. This result in consistent with the previously observed primary role of pelvis displacement in predicting step width, and only a secondary role of other biomechanical factors [Stimpson et al. 2018]. Equations 2, 3, and 4 all produced significant increases in step start paretic $\rho_{disp}$ relative to the baseline walking condition. Based on these results, we chose to apply Equation 2 in this study's main experiment. This choice was based on our ultimate goal of developing a tool that can be implemented in a clinical setting. Equation 2 involves only displacement-dependent forces, which could be feasibly implemented using a passive system of pulleys and springs (which can be simplistically thought of as converting displacements to forces). The inclusion of a velocity term in Equations 3 and 4 would require a more complex design, possibly involving damping (or negative damping) elements.

For Equation 2, we calculated an effect size of 1.2 for our primary outcome measure of step start paretic $\rho_{disp}$, as assistance caused an increase in this measure of 0.17±0.14 (mean±SD). Given this effect size, we calculated that a sample size of 10 would be required to achieve 80% power with an alpha value of 0.025 – as planned for our main experiment.

***Appendix References***


Browning RC, Kram R. Effects of obesity on the biomechanics of walking at different speeds. Med Sci Sport Exerc. 2007;39:1632-1641.

Bruijn SM, van Dieen JH. Control of human gait stability through foot placement. J R Soc Interface. 2018;15:20170816.

Hak L, Houdijk H, Steenbrink F, Mert A, van der Wurff, Beek PJ, van Dieen JH. Speeding up or slowing down? Gait adaptations to preserve gait stability in response to balance perturbations. Gait Posture. 2012;36:260-264.

Heitkamp LN, Stimpson KH, Dean JC. Application of a novel force-field to manipulate the relationship between pelvis motion and step width in human walking. IEEE Trans Neur Syst Rehabil Eng. 2019;27:2051-2058.

Nyberg ET, Broadway J, Finetto C, Dean JC. A novel elastic force-field to influence mediolateral foot placement during walking. IEEE Trans Neur Syst Rehabil Eng. 2017;25:1481-1488.

Rankin BL, Buffo SK, Dean JC. A neuromechanical strategy for mediolateral foot placement in walking humans. J Neurophysiol. 2014;112:374-383.

Stimpson KH, Heitkamp LN, Horne JS, Dean JC. Effects of walking speed on the step-by-step control of step width. J Biomech. 2018;68:78-83.

Stimpson KH, Heitkamp LN, Embry AE, Dean JC. Post-stroke deficits in the step-by-step control of paretic step width. Gait Posture. 2019;70:136-140.

Wang Y, Srinivasan M. Stepping in the direction of the fall: the next foot placement can be predicted from current upper body state in steady-state walking. Biol Lett. 2014;10:20140405.

***Appendix Tables and Figures***

| Participant # | Age (yr) | Time since stroke (mo) | Paretic side | Gender | LE FM score | Walking speed (m/s) |
|---|---|---|---|---|---|---|
| 1 | 53 | 19 | L | F | 24 | 0.3 |
| 2 | 23 | 74 | R | M | 24 | 0.8 |
| 3 | 71 | 39 | R | M | 23 | 0.4 |
| 4 | 82 | 27 | R | F | 26 | 0.25 |
| 5 | 33 | 103 | L | F | 21 | 1.2 |
| 6 | 38 | 80 | L | M | 15 | 0.45 |
| 7 | 71 | 59 | R | M | 30 | 0.3 |
| 8 | 76 | 13 | R | F | 34 | 0.7 |
| 9 | 54 | 26 | L | F | 24 | 0.7 |
| 10 | 56 | 201 | R | F | 34 | 0.55 |
| 11 | 52 | 36 | L | F | 24 | 0.6 |
| 12 | 76 | 25 | R | M | 25 | 0.3 |
| *Group summary* | 57±19 | 59±53 | 5L / 7R | 7F / 5M | 25±5 | 0.55±0.28 |

Table S1. Participant demographics, clinical measures, and treadmill walking speeds.

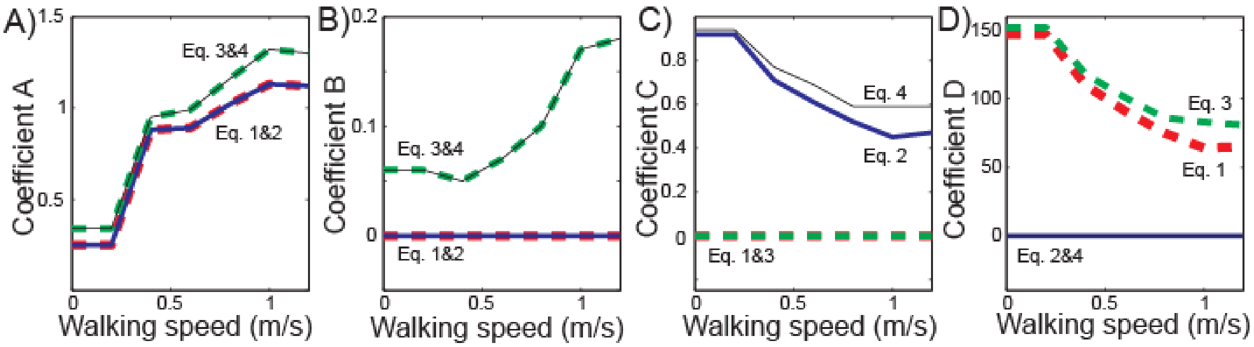


Figure S1. Force-field coefficient values are illustrated for the four candidate equations, with each coefficient (A-D) corresponding to the panel label. These values vary with walking speed, based on empirically-derived best-fit coefficients from neurologically-intact control participants walking at a range of speeds, and interpolated between these speeds. The four equations are indicated with labels (Eq. 1 through Eq. 4) on each panel.

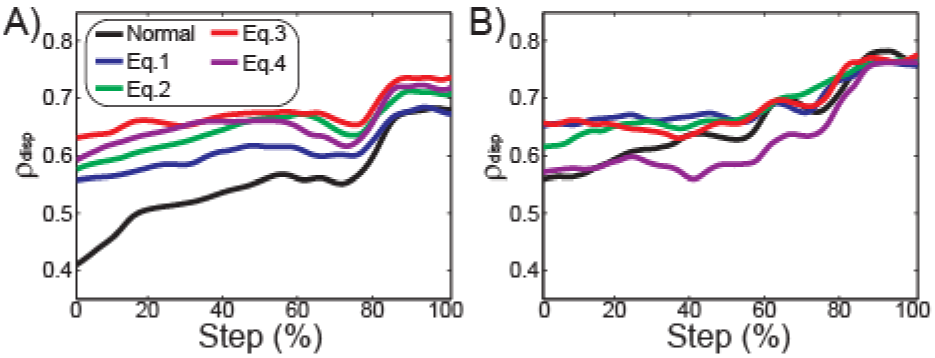


Figure S2. $\rho_{disp}$ values calculated throughout the step, from step start (0%) to step end (100%). The direct effects and after-effects of each assistance equation are illustrated for steps taken with the paretic leg (A) and the non-paretic leg (B). Plots illustrate the mean value of these metrics across participants.

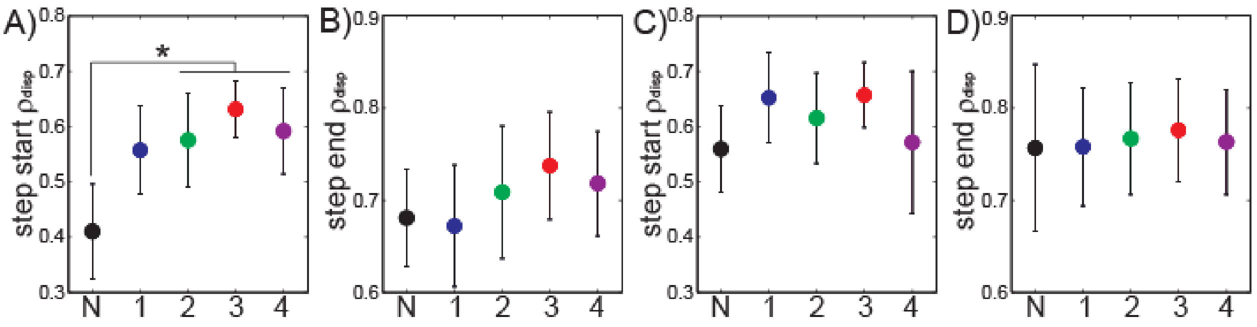


Figure S3. Force-field effects on $\rho_{disp}$ values. Each panel illustrates a comparison between Normal walking (abbreviated N on x-axis) and the four candidate assistive equations (indicated by numbers 1-4). These comparisons were performed for step start paretic $\rho_{disp}$ (A), step end paretic $\rho_{disp}$ (B); step start non-paretic $\rho_{disp}$ (C); and step end non-paretic $\rho_{disp}$ (D). Dots indicate group means, and error bars indicate 95% confidence intervals. The asterisk indicates a statistically significant post-hoc difference between conditions.
